# Ish: SIMD and GPU Accelerated Local and Semi-Global Alignment as a CLI Filtering Tool

**DOI:** 10.1101/2025.06.04.657890

**Authors:** Seth Stadick

## Abstract

**Background:** Filtering records using command line tools is a staple of Bioinformatics. In analysis pipelines and in day-to-day research tools such as awk, grep, and cut are the workhorses of much of our data crunching. To date, there is no command line utility for performing index-free alignment-based filtering of records.

**Results:** Ish is a classic and composable unix-style CLI tool. It takes a query and filters the input target records to only those that match the input query with a threshold alignment score, using a selectable alignment algorithm. The core alignment algorithms for Ish meet or exceed the performance of their reference implementations for both SIMD and GPU alignment, as measured by gigacell updates per second (GCUPs).

**Conclusions:** Ish fills a longstanding gap in Bioinformatics tooling by offering a CLI utility for index-free alignment-based record filtering. Additionally, it provides meaningfully improved versions of the classic striped local and semi-global full dynamic programming algorithms, along with a GPU-based semi-global alignment method.

## 1 Background

To date, there are very few index-free approximate match CLI tools. There are many highly specialized tools in the field of bioinformatics to perform approximate matching (aka alignment) on different sequence types, but all require the preparation of reference sequences into a bespoke index format ahead of time (see BWA-MEM [1], for example). In addition, they are usually specific to either protein or nucleotide alignments. Outside of bioinformatics, there are even fewer tools available. The most notable tool is agrep (from TRE [2]), an approximate matching regular expression implementation that can allow for up to N mismatches and specifying mismatch, insertion, and deletion scores. TRE is single-threaded and does not support common record formats in bioinformatics (FASTA, FASTQ, SAM).

Sequence search and filtering are a deep field of research because of the pervasiveness and usefulness of the problem [3]. Sequences are composed of an alphabet of symbols that can be composed in any order and at any length. Sequence search comes in multiple forms, each of which has a highly divergent set of optimizations and solution spaces:

- Exact Matching: Find an occurrence of a query sequence in a target.
- Pattern Match: Use a symbolic pattern (i.e. regex) to find an occurrence of that pattern in the target.
- Approximate Matching: Find a subsequence that is most similar to the query (based on a similarity metric) in the target.

Ish uses optimal alignment algorithms with adaptive gap scoring (affine costs), specifically, local alignment and semi-global alignment. The implementation uses the striped SIMD algorithm described by [4], and refined by SSW [5], with Parasail [6] as the reference point, with some modifications (see Implementation 2).

The core of the striped SIMD algorithm is the query profile, reviewed in Figure 1. The query profile precomputes the cost vectors of the query for each alphabet character. This allows for efficient comparison of the query against the reference, obviating the need to lookup the value in the scoring matrix. For an in-depth review of the striped SIMD algorithm, see [4, 5, 7]

**Fig. 1.**
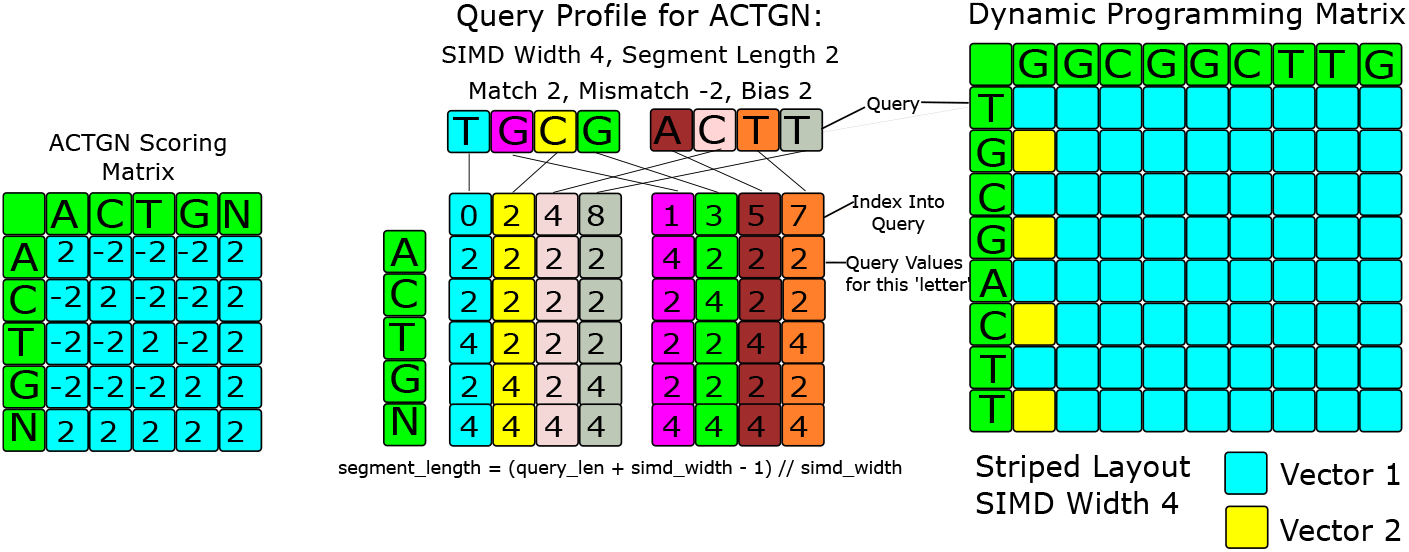
Striped SIMD Query Profile. The query profile is a list of SIMD vectors that pre-loads the score for each query letter to avoid lookups. The middle matrix here is the query profile. Each row is the pre-loaded scores between the query and the given nucleotide. The split in the middle represents the two SIMD vectors needed to represent the length eight query, and how the striping works with the first vector for alphabet character A uses the “striped” pattern of 0, 2, 4, 6, and the second vector covers 1, 3, 5, 7. The dynamic programming matrix can than be iterated over in a striped fashion, simd width query letters are compared against the current target letter at once. [4]

**Fig. 2.**
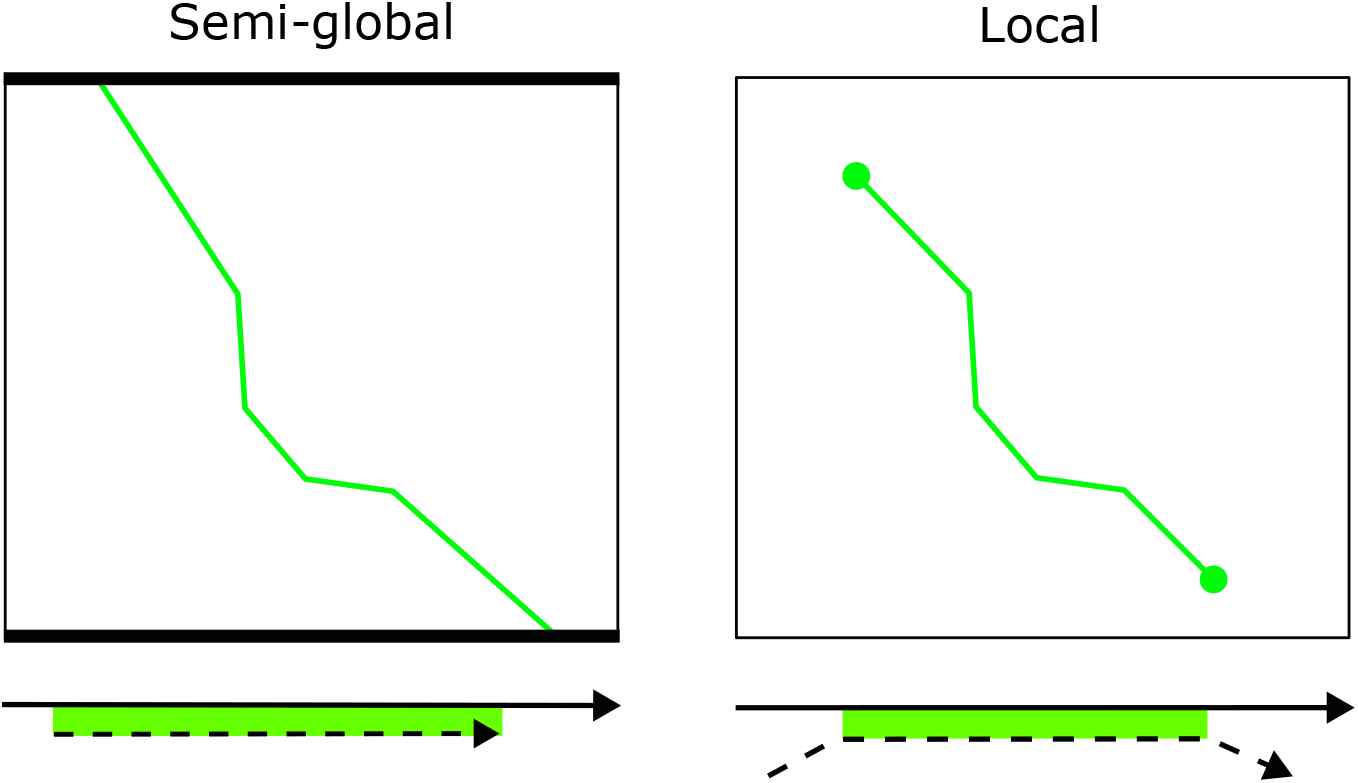
Visualization of Semi-global and Local Alignment Methods. Each rectangle is a dynamic programming matrix silhouette, and the green lines are paths through the matrix representing the alignment. The dark bars along the perimeter indicate where the alignment can start and end. The arrows with green highlighting represent the sequence alignments in the matrices above [3]

GPU-based alignment is still a developing area. There are two strategies used on the GPU that have been picked up by the CPU. Intra-task parallelization is parallelization of the processing of a single pairwise alignment [7]. Inter-task parallelization is the parallelization of many pairwise alignments at once. Striped SIMD is an example of Intra-task parallelization. Both approaches have been attempted on the GPU. Often a combination of two is used to maximize performance. SIMD or SIMT for intra-task parallelization, and multithreading (CPU or GPU) for inter-task parellelization. For example, the current fastest local alignment method on GPUs is CUDASW which uses warp-based SIMT and efficient tiling to achieve orders of magnitude better performance than any other implementation [8].

## 2 Implementation

### 2.1 Mojo

Ish is implemented in Mojo [9], which was first released in 2023 [10]. Mojo is focused on high-performance computing across heterogeneous accelerators, specifically GPUs. Syntactically, Mojo is part of the Python family. Memory management in Mojo is similar to that in Rust, using an ownership system which prevents errors such as use-after-free, double-free, and memory leaks. Mojo was chosen for three reasons: to demonstrate its usage in a bioinformatics tool, its support for SIMD programming, and its support for GPU programming [11]. In all three categories, it excelled, and each will be discussed in greater detail in subsequent sections. Version 25.2 was used for all the benchmarks shown here.

### 2.2 Command Line Interface

#### 2.2.1 Selectable Record Type

Ish is a command line tool developed with bioinformatics applications in mind but is broadly applicable for approximate matching uses across disciplines. Version 1.0 supports FASTA, and Line record types, allowing it to align only against the sequences of interest in those record types. In the case of FASTA this results in a substantial performance improvement, since Ish is able to do less work than a tool like agrep by not attempting to match against header lines. Additionally, being record type aware allows Ish to output records in the correct format, so a matched FASTA record will output the header and sequence instead of just the matched line.

Allowing for matching against lines as a record type allows Ish to be used in contexts outside of bioinformatics, such as searching a codebase, regular text search, etc. By default, ish will recursively search all non-dotfile non-binary files in your current directory (or selected directory) similar to ripgrep [12], and output matched lines with the filepath and line number formatted correctly as terminal links [13].

### 2.2.2 Selectable Alignment Method

Ish supports two different alignment methods, local and semi-global. Local was selected because it allows for finding the most homologous subsequence between two sequences [3], which is useful for splice site detection [14, 15], conserved domain detection [16], etc. Additionally, it has useful properties for working with SIMD, such as a minimum score of 0 which can increase performance by removing the need to perform an underflow check.

Semi-global was selected because it generalizes to most other forms of alignment by allowing one to specify which combination ends of the query and target you would like to be “free” from a scoring perspective [3]. By default, ish allows for both ends of the query sequence to be free but allows specifying any combination.

#### 2.2.3 Selectable Scoring Matrices

Scoring matrices play an important role in producing high-quality alignments [17] by allowing the alignment to take biology into account. In ish we support a generic ACTGN scoring matrix for nucleotides, Blosum62 [17] for protein sequences, and ASCII for everything else.

In addition to alphabet-specific scoring matrices, ish supports affine gap penalties across all alignment methods, further improving alignments by allowing them to represent biology more closely, in the case of protein and nucleotide sequences [18].

### 2.3 Striped SIMD Alignments

In addition to porting the Parasail implementations to Mojo, we provide several novel improvements and advancements.

ish supports AVX512 instructions for both local and semi-global alignments, which are not supported in other striped-simd alignment methods that were reviewed. AVX512 vectors are twice as large space as AVX2 vectors, allowing for even greater parallelism, which is most important for very long queries. Although not explicitly demonstrated here, AVX512 offers a more complete set of instructions that can reduce operations that previously needed multiple instructions to a single call [19, 20].

A particularly interesting finding, which is directly a side effect of using Mojo’s SIMD abstraction, was the usage of SIMD strip mining (unrolling, or otherwise using multiple simd vectors at once) on aarch64 to mimic AVX2 [21]. If the SIMD width is configured to be a multiple of the hardware SIMD width, the Mojo compiler will act by simulating larger SIMD vectors and generate code that has a performance profile similar to the larger hardware vector size; see Listing 1. It is not as performant as the actual AVX2 registers, for example, but it is substantially more performant on long queries than the vector size of the base hardware (see Results 3). While the example shown is rather simplistic, the real advantage is when there are interlane dependencies or reductions. The compiler is able to handle these scenarios seamlessly and allows one to try wider or smaller widths.

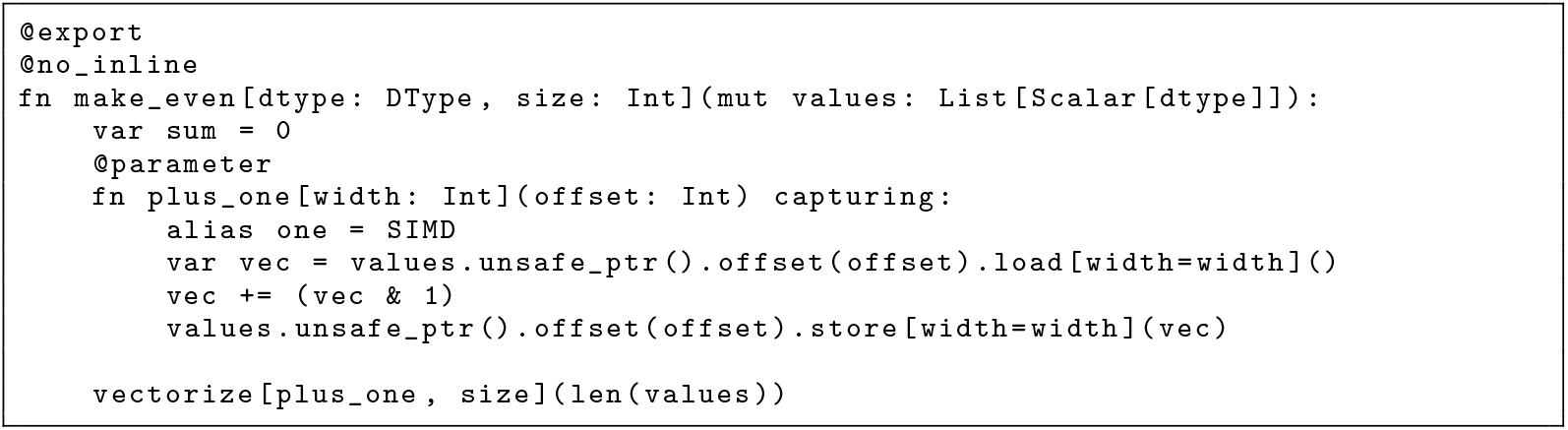
Listing 1 Vectorized Function. Function to convert all array values to even numbers.

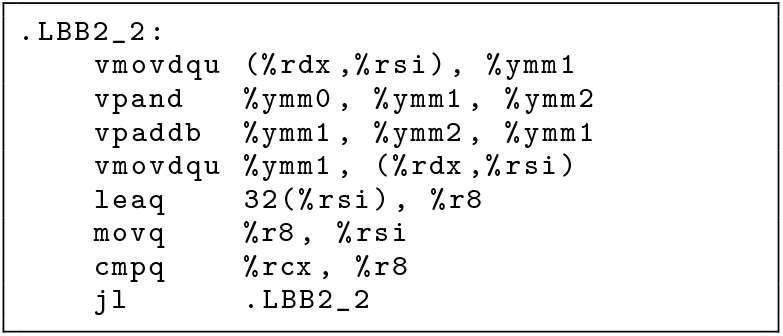
Listing 2 No Unrolling. Inner Loop for AVX2 When Width Equals Harware Width

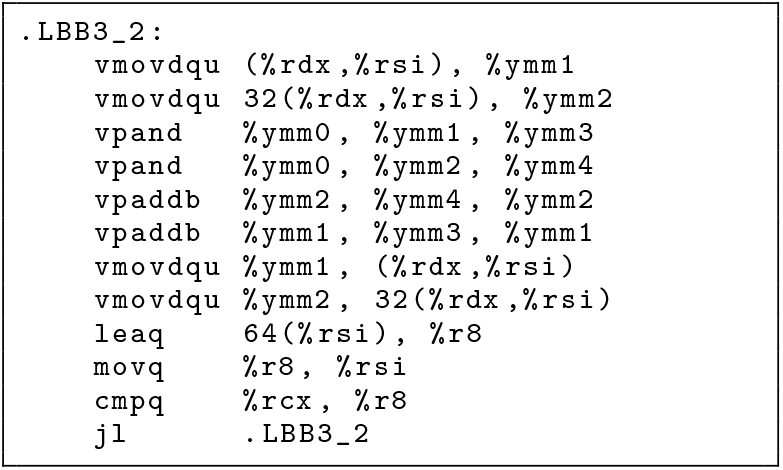
Listing 3 With Unrolling. Inner Loop for AVX2 When Width Equals 2x Harware Width

In the striped-semi-global implementation we also made several improvements over the reference Parasail implementations. To implement striped-semi-global alignment, we needed to create our own SIMD saturating add and subtract in Mojo. However, checking for overflow and underflow on signed integers is very expensive compared to unsigned integers [22] because there is no defined behavior for overflow or underflow of signed integers, and therefore an explicit check has to be made, which necessitates both widening the integer to the next power of two width, as well as introducing a branch in the code path.

To account for this shortcoming, we use unsigned integers, but shift them up to their midpoint and set the midpoint as “0”, allowing for the same range of values as a signed integer. This allows for eliding half the bounds checks by adjusting the saturation checkpoint to the minimum of the unsigned integer max minus the absolute value of the largest negative score and minus the largest positive score, respectively. When a negative score is subtracted, it can then cause intentional underflow, into the positive high range (see the Parasail [6] implementation for a detailed description of the semi-global implementation). See Figure 3. This does have the downside of marginally reducing the range of positive scores, but such an effect should be negligible for most common scoring matrices.

**Fig. 3.**
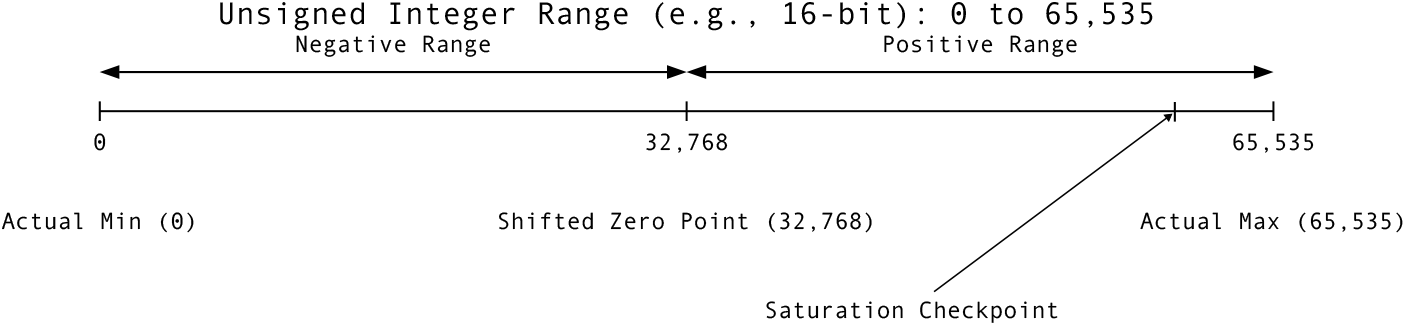
Saturation Check Elision. By using an unsigned integer for scoring and shifting the “zero” value to be the midpoint of that integer, wrapping underflow can be relied upon to put a value in to the high range. As such, a single saturation checkpoint value can be used to check if overflow or underflow may happen by setting the checkpoint to the minimum of the integer max minus the absolute value of the largest negative score, and minus the largest positive score, respectively.

### 2.4 GPU Alignment

The approach taken for doing GPU based alignment is intentionally naive and runs a scalar version of semi-global alignment (from Parasail [6]) on N threads of the GPU, with a definable coarse-graining factor. The method moves the query to the outer loop and the reference to the inner loop, which allows for a better likely-case memory usage where the query is smaller than the target. Additionally, we cap the target lengths that go to the GPU to a fixed amount (see GPU / CPU Parallelization Dataflow 2.5.2). Capping the length ensures that each warp should have approximately the same amount of work to do [23].

Although our algorithm performs reasonably fast, it is still more efficient to use the CPU on smaller inputs. On inputs where the configurable batch size is roughly three times smaller than the file size, the overhead of setting up the GPU buffers becomes worthwhile. When multiple GPUs are detected, the work will split between them. The size of the input buffer is multiplied by the number of available GPUs, and then the work is evenly split between them.

Lastly, given the necessity of using fixed size buffers on the GPU, we have parameterized the max query length and the max target length into subsets of likely buckets. This ensures that buffers will be close to right-sized for any input query or discovered max target length. In the event that the lengths are outside what is supported, Ish falls back to CPU processing.

### 2.5 Data Flow

#### 2.5.1 CPU Parallelization

The concurrency model in Mojo v25.2 does not have a mechanism to do any work in a way that does not block the main thread outside of offloading to the GPU. The only concurrency available is through a standard library function called parallelize, which takes a closure and a number of works and launches parallel tasks for you. This results in a blocky form of parallelism in which the work is built up serially, processed in parallel, and then output serially. This could be streamlined by the addition of threads or an async runtime.

Figure 4 demonstrates the end-to-end CPU workflow. Records are accumulated in batches before they are sent for parallel processing. N threads are created to process the input, and each thread is assigned a linear set of input records to process. Once processed, the records result is placed in the pre-allocated output buffer matching index position. Upon completion, the matches are returned to the main loop and written.

**Fig. 4.**
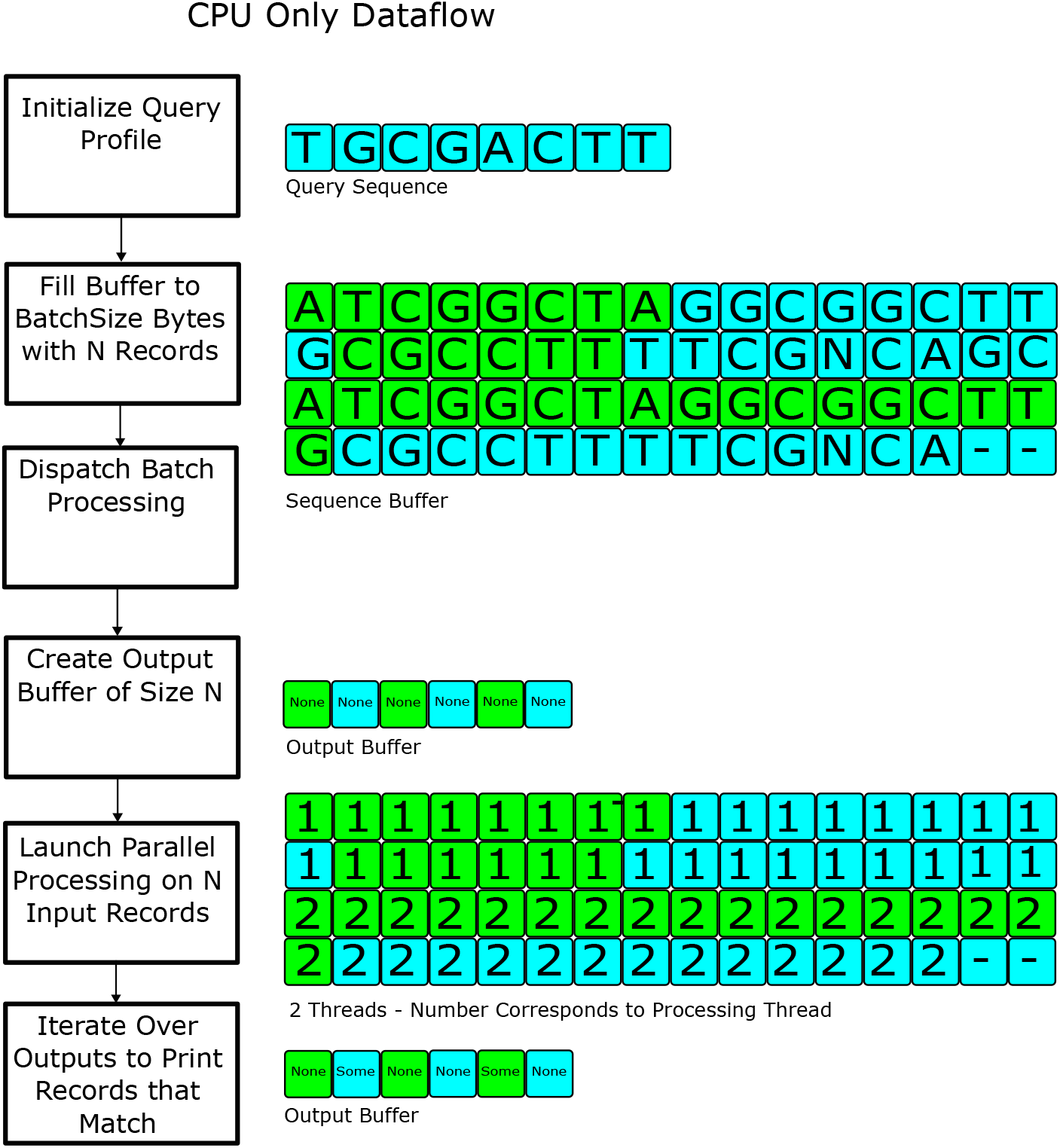
CPU Dataflow. Records are accumulated as a batch of configurable size as they are read from input files. Once the batch size is reached, or end-of-file is hit, the batch is dispatched for processing. The processing step pre-allocates the result locations, and then launches threads (as many as specified via the CLI, or available) to process the data. In the example above 2 threads are launched and the work is split such that each thread can operate on continuous chunks of the input batch. Each records result is written to its corresponding index in the output buffer. Finally, the outputs are iterated over and any records that have a result are written. The blue and gree segments indicate different records.

**Fig. 5.**
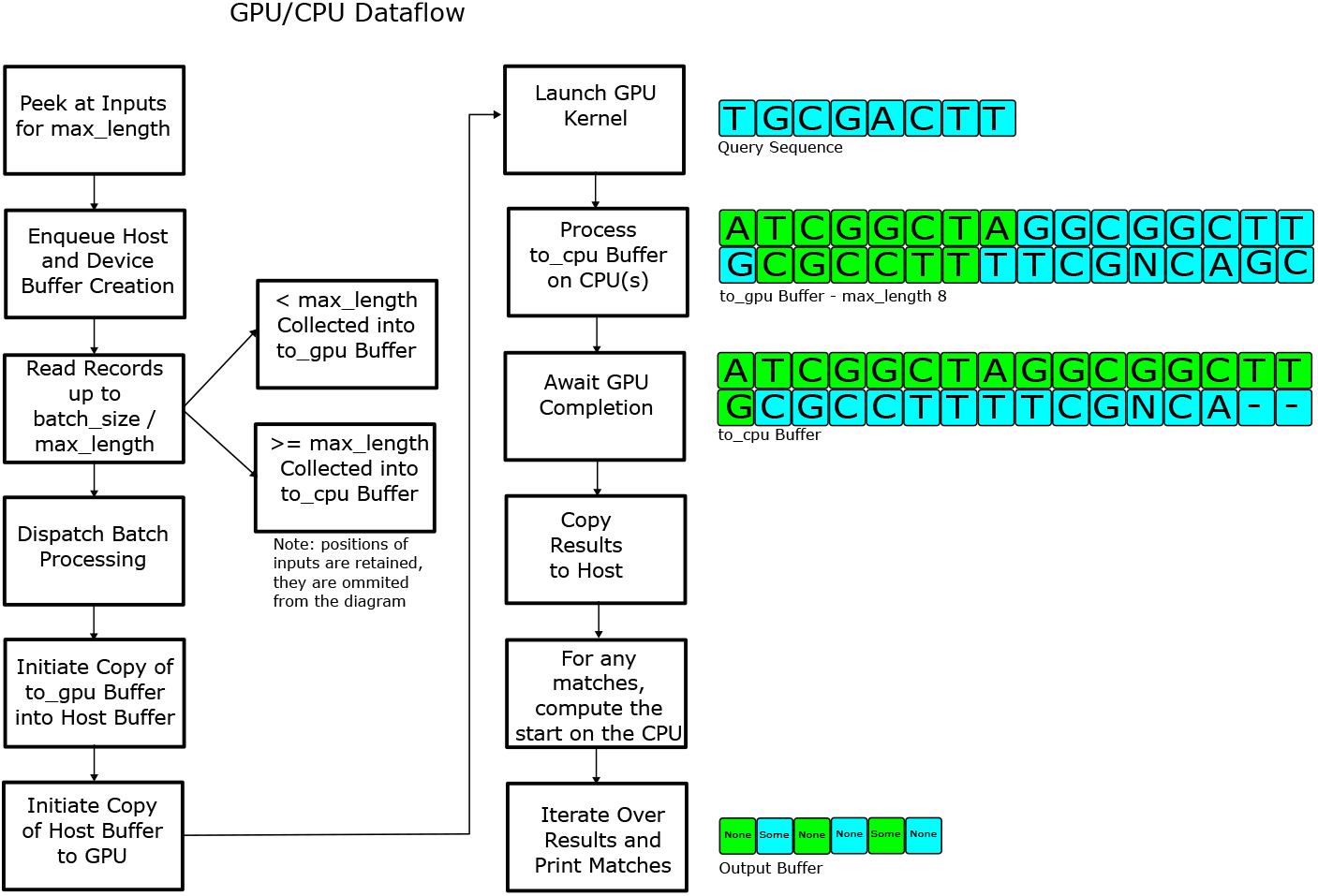
GPU Dataflow. First, input is first checked to determine a good max length buffer to use for the GPU buffers. Host-side and GPU-side buffers are allocated based on the chosen max length. Input records are collected into the GPU and CPU based buffers based on their lengths. The data transfers, and kernel execution are enqueued on the GPU and the CPU records are processed during the GPU execution. Results are then iterated over and written serially.

#### 2.5.2 Hybrid GPU / CPU Parallelization

Before accumulating records into batches, the input is first checked to determine a good max length buffer to use by reading the first several thousand bytes of the input; this allows for right-sizing the inputs to the GPU which depend on the max length being close to the median length of the inputs.

The CPU (host) side and GPU side buffers are allocated based on the chosen max length. The input records are parsed and collected into two buckets: those records that are less than or equal to max length, destined for the GPU, and those that are greater than max length, which are destined for the CPU. When a GPU batch is filled, the CPU and GPU buffers are dispatched for processing.

The copy of the GPU records to the GPU is initiated, followed by the launch of the GPU kernel. While those functions are running asynchronously, the CPU records are processed in the same manner as described in the CPU parallelization data flow. Upon completion of the CPU processing, the GPU functions are awaited and complete with a copy of the results from the GPU to the host. The results are then iterated over and written in serial order.

## 3 Results

### 3.1 Striped SIMD Alignment

The following benchmarks reproduce a subset of those performed by Rognes [24] and Daily [6]. The ish striped SIMD implementations were compared against Parasail’s using Parasail’s aligner tool that is meant for benchmarking. A similar aligner tool was created for ish that benchmarks just the alignment method and output generation.

All alignment results were validated against Parasail’s outputs to ensure correctness. The speed is reported in billion (giga) cell updates per second (GCUPS), where a cell is a value in the dynamic programming matrix. The loading time and profile creation time were excluded from these results, and all results in this section are for a single thread. The subset of sequences used are listed in Table 1, and the query sequences were the same as the Parasail manuscript and had lengths ranging from 24 to 5478. Only the Blosum62 [17] scoring matrix was used since that is what Daily used for comparison between tools. Unlike the original Parasail manuscript, a gap open of 3 and a gap extend of 1 was used. Parasail v2.6.2 was used for all testing and was compiled from source for each machine following the build procedure in the README.

**Table 1.**
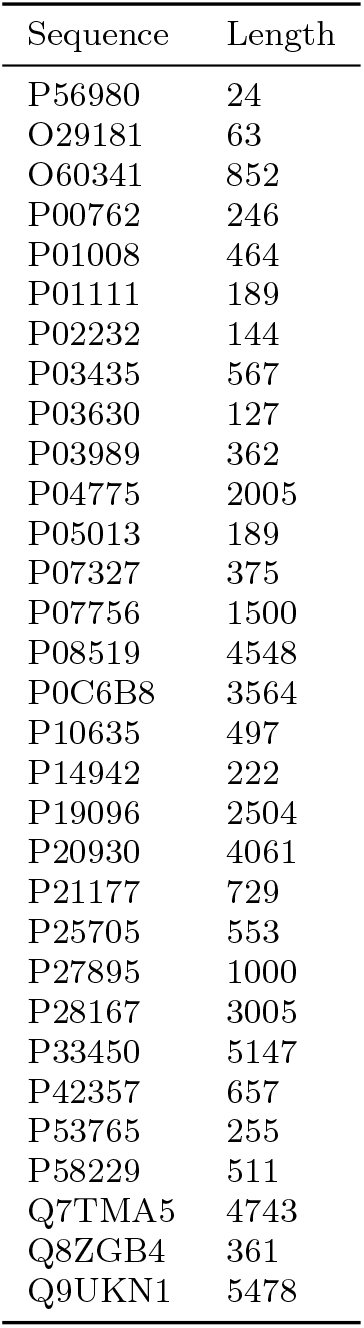
UniProt Accession Numbers. Query sequences used, and their associated lengths. This is a subset of the query sequences used in the Parasail manuscript.

| Sequence | Length |
| --- | --- |
| P56980 | 24 |
| O29181 | 63 |
| O60341 | 852 |
| P00762 | 246 |
| P01008 | 464 |
| P01111 | 189 |
| P02232 | 144 |
| P03435 | 567 |
| P03630 | 127 |
| P03989 | 362 |
| P04775 | 2005 |
| P05013 | 189 |
| P07327 | 375 |
| P07756 | 1500 |
| P08519 | 4548 |
| P0C6B8 | 3564 |
| P10635 | 497 |
| P14942 | 222 |
| P19096 | 2504 |
| P20930 | 4061 |
| P21177 | 729 |
| P25705 | 553 |
| P27895 | 1000 |
| P28167 | 3005 |
| P33450 | 5147 |
| P42357 | 657 |
| P53765 | 255 |
| P58229 | 511 |
| Q7TMA5 | 4743 |
| Q8ZGB4 | 361 |
| Q9UKN1 | 5478 |

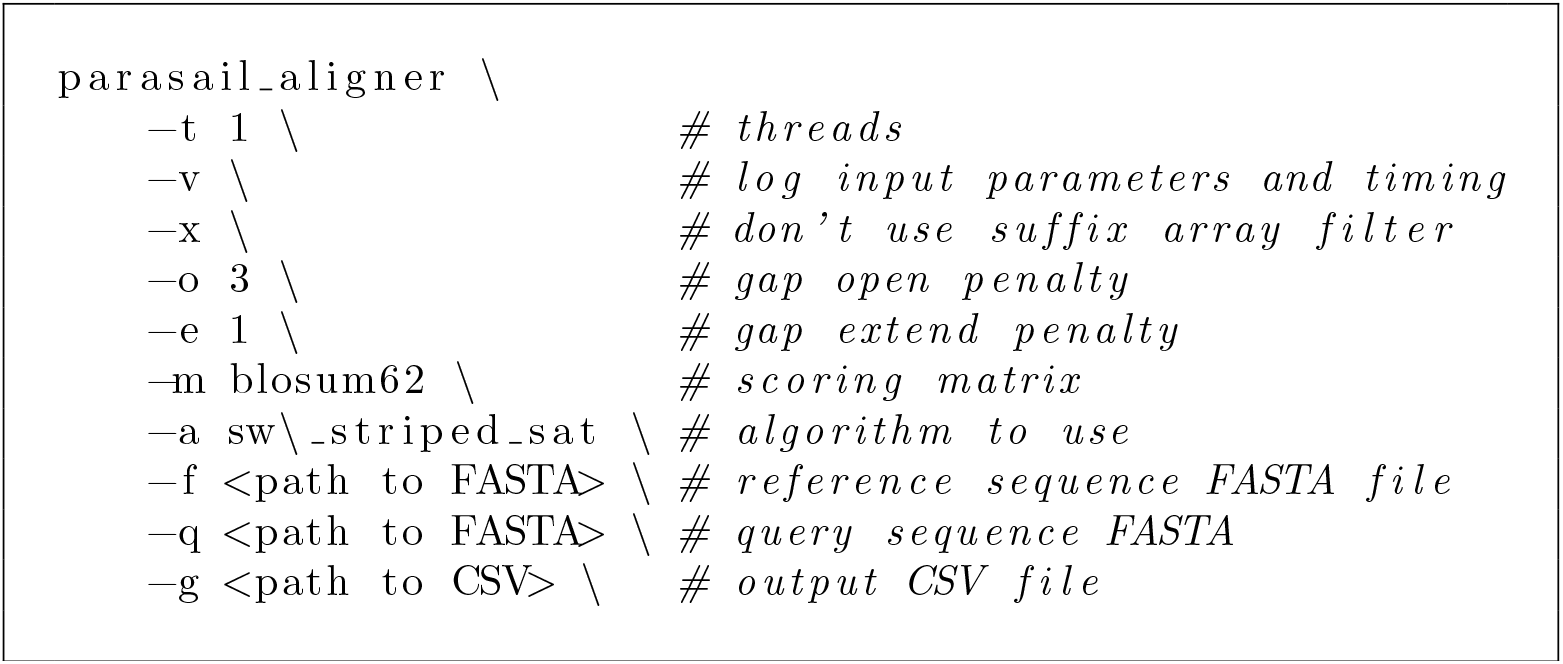
Listing 4 CLI Invocation for Parasail. CLI options used for running parasail aligner.

#### 3.1.1 Local Alignment Evaluation

Figure 6 and Table 2 3 shows the performance of Parasail and Ish over the full range of query sequence lengths using the adaptive scoring mechanism, which is the most common for local alignment. The adaptive mechanism will begin by using u8s to hold scores, and when an overflow is detected, switch to u16s (see SSW documentation for further discussion [5]. Furthermore, Figure 6 shows the different SIMD registers used.

**Table 2.** x64 Local Alignment Comparison. Average GCUPs per query length bin comparing Ish and Parasail and showing the % difference relative to Parasail. A positive percent is an improvement where a negative would be a regression. 64/32 widths not included since there is no comparison point with Parasail.

| Query Length | Width Config | ISH GCUPS | Parasail GCUPS | % Difference |
| --- | --- | --- | --- | --- |
| 0-200 | 16/8 | 2.11 | 2.11 | -0.2% |
| 200-400 | 16/8 | 3.06 | 3.00 | 2.1% |
| 400-600 | 16/8 | 3.45 | 3.35 | 2.9% |
| 600+ | 16/8 | 3.93 | 3.82 | 2.9% |
| 0-200 | 32/16 | 2.65 | 2.53 | 4.8% |
| 200-400 | 32/16 | 4.56 | 4.10 | 11.3% |
| 400-600 | 32/16 | 5.65 | 4.87 | 15.8% |
| 600+ | 32/16 | 7.48 | 6.16 | 21.4% |

**Table 3.** aarch64 Local Alignment Comparison. Average GCUPs per query length bin comparing Ish and Parasail and showing the % difference relative to Parasail. A positive percent is an improvement where a negative would be a regression. 32/16 widths not included since there is no comparison point for Parasail.

| Query Length | Width Config | ISH GCUPS | Parasail GCUPS | % Diffence |
| --- | --- | --- | --- | --- |
| 0-200 | 16/8 | 1.67 | 1.43 | 16.8% |
| 200-400 | 16/8 | 2.26 | 1.99 | 13.4% |
| 400-600 | 16/8 | 2.47 | 2.23 | 11.2% |
| 600+ | 16/8 | 2.70 | 2.59 | 4.5% |

**Fig. 6.**
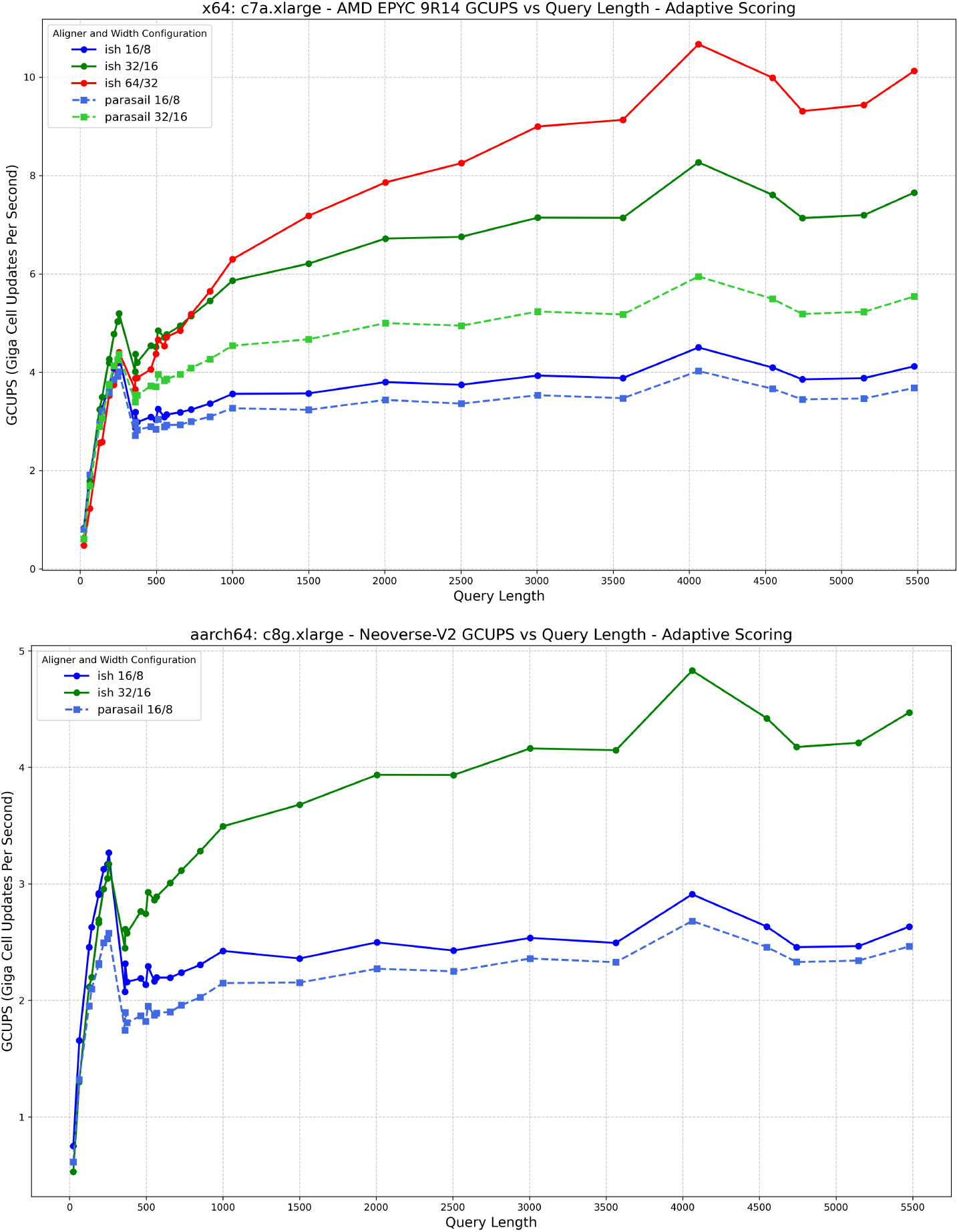
Striped SIMD Local Alignment Comparison. Comparison of Striped SIMD local alignment methods between Parasail and with different SIMD widths over a range of input query lengths. The top chart is for x64 and the bottom chart is for aarch64, which each offer different SIMD instruction sets and capabilities. Ish results are shown as solid lines and Parasail as dashed lines. The SIMD widths represent the vector width for an 8 bit container, and 16 bit container, respectively. Notable here is the over-sizing of the vectors for ish for aarch64, which improves performance on long queries.

The performance of ish is in line with or slightly improved when compared to the standard set by Parasail. Notably, the larger vector sizes usable by Ish perform better on long query sequences than Parasail. On x64 ish takes advantage of AVX512 instructions, which are slower at short lengths but outstrip AVX2 with queries of approximately length 750 or greater.

On aarch64 benchmarks were added to test the Mojo compiler’s ability to generate SIMD strip-mining optimizations by increasing the SIMD width to a multiple (2) of the hardware vector size. When doing this, on long queries, ish is able to somewhat mimic the effect of having larger hardware registers. Using widths 32/16 when the hardware supports 16/8 does not quite double performance in the same way that going from SSE to AVX2 does, but it still provides a substantial boost for long queries.

#### 3.1.2 Semi-Global Alignment Evaluation

Figure 7 and Tables 4 5 show the performance of Parasail and Ish across the full range of query sequence lengths using the word (16 bit containers) scoring mechanism. Byte size scoring was skipped since 8 bit containers overflow quickly with semi-global alignment scoring, as noted in the Parasail manuscript.

**Table 4.** x64 Semi-Global Alignment Comparison. Average GCUPs per query length bin comparing Ish and Parasail and showing the % difference relative to Parasail. A positive percent is an improvement where a negative would be a regression.

| Query Length | Width Config | Ish GCUPS | Parasail GCUPS | % Difference |
| --- | --- | --- | --- | --- |
| 0-200 | 16/8 | 2.17 | 1.86 | 16.8% |
| 200-400 | 16/8 | 3.17 | 2.63 | 20.8% |
| 400-600 | 16/8 | 3.62 | 2.95 | 22.4% |
| 600+ | 16/8 | 4.18 | 3.43 | 21.7% |
| 0-200 | 32/16 | 2.70 | 2.22 | 21.6% |
| 200-400 | 32/16 | 4.64 | 3.61 | 28.4% |
| 400-600 | 32/16 | 5.70 | 4.36 | 31.0% |
| 600+ | 32/16 | 7.55 | 5.59 | 35.1% |

**Table 5.**
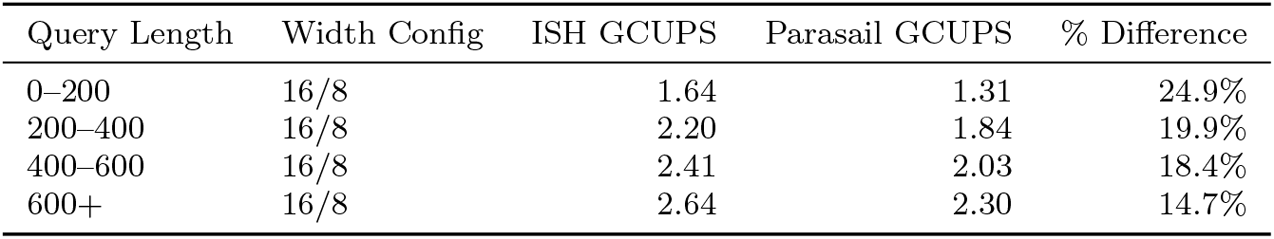
aarch64 Semi-Global Alignment Comparison. Average GCUPs per query length bin comparing Ish and Parasail and showing the % difference relative to Parasail. A positive percent is an improvement where a negative would be a regression.

**Table 6.** Searching Line Records.

| Tool | Query Length | Time | Times Slower |
| --- | --- | --- | --- |
| TRE agrep 1 thread | 50 | 371.480 s | 75.18 |
| ish cpu 1 thread | 50 | 124.641 s | 25.23 |
| ish cpu 24 threads | 50 | 9.505 s | 1.92 |
| ish 1 gpu | 50 | 5.000 s | 1.01 |
| ish 2 gpu | 50 | 4.941 s | 1.00 |

Comparison of line-based searches between agrep and ish. agrep is single threaded and allows up to 12 mismatches where ish uses a score threshold of .75 (alignment score must be at least 75% of the maximal alignment score for that query). Times Slower is relative to the fastest time in the benchmark group.

**Table 7.** FASTA vs Line Records.

| Tool | Query Length | Time | Times Slower |
| --- | --- | --- | --- |
| TRE agrep 1 thread | 50 | 371.480 s | 92.96 |
| ish cpu 1 thread | 50 | 81.412 s | 20.37 |
| ish cpu 24 threads | 50 | 7.181 s | 1.80 |
| ish 1 gpu | 50 | 3.996 s | 1.00 |
| ish 2 gpu | 50 | 4.108 s | 1.03 |

Comparison of searching FASTA records vs searching a FASTA file as lines. agrep uses line based searching. All runs of ish presented here use FASTA record searching. Times Slower is relative to the fastest time in the benchmark group.

**Fig. 7.**
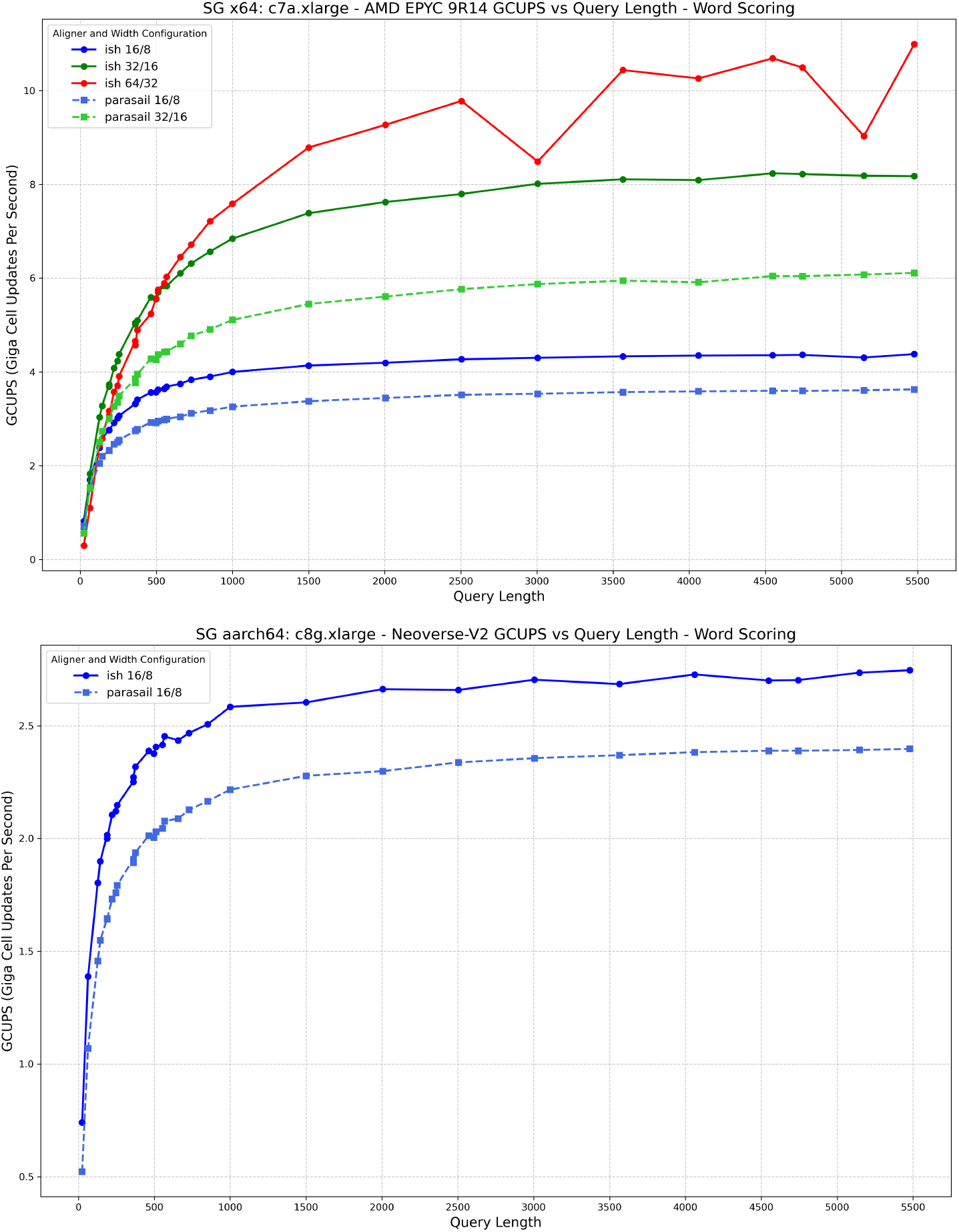
Striped SIMD Semi-Global Alignment Comparison. Comparison of Striped SIMD semi-global alignment methods between Parasail and with different SIMD widths over a range of input query lengths. The top chart is for x64 and the bottom chart is for aarch64, which each offer different SIMD instruction sets and capabilities. Ish results are shown as solid lines and Parasail as dashed lines. Only 16 bit containers (also referred to as word scoring) was used for this benchmark.

The performance of ish is again in line with or slightly improved compared to the standard set by Parasail. The improvement is reflective of the algorithm change to make use of unsigned integers and elide a lower bounds check, as described in the Implementation section.

### 3.2 GPU Alignment

Figure 8 illustrates the raw performance of the GPU kernel used in ish. Specifically, it measures the GCUPs starting from the start of the data copy to the GPU, and stopping after the transfer off the GPU. Input sequences are sorted by length and per-GPU. The performance within Ish is not directly used because the overhead of parsing FASTA records is too high to properly utilize the throughput of the GPU. The same benchmarking tool was used for GPU alignments as was used for SIMD alignments, structured after the Parasail aligner.

**Fig. 8.**
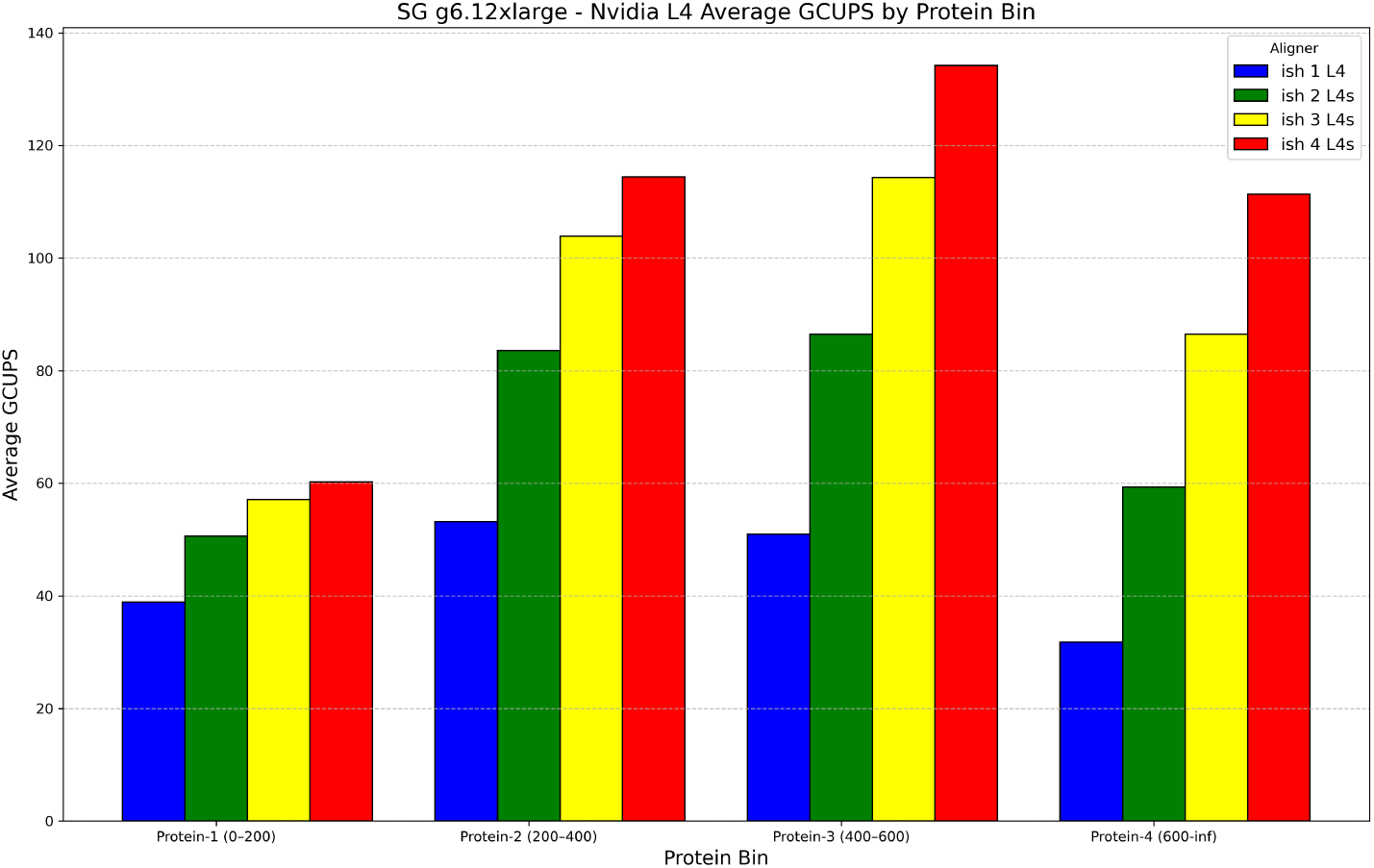
GPU Based Semi-Global Alignment Performance. Semi-Global Alignment performance in GCUPs binned by query length. Each color represents the number of GPUs used, demonstrating ish’s ability to scale with multiple GPUs.

ADEPT [25] provides the best comparison for Ish. Measurements are broken up as in the ADEPT manuscript. ADEPT uses a cumulative mean for each bin (they use 0-200, 0-400, 0-600) where this benchmark uses discrete bins. ADEPT was not run for comparison, but their manuscript has comparable GCUP values. In particular, ADEPT was benchmarked on an older generation of hardware than Ish. Broadly, Ish and ADEPT have comparable performance on a single GPU, with ADEPT seeming to have better multi-gpu support. Generally, Ish performs marginally better on medium-to-long query sequences as well where ADEPT excels at shorter sequences. The same data set and scoring as the SIMD benchmarks was used, but scaled up by 5x to better utilize the GPU compute.

### 3.3 CLI Tool Comparisons

The only CLI tool that works like ish, by fuzzy matching input records and filtering them based on a score, is agrep [2]. agrep works on line-based records by default but does allow for defining a custom record delimiter regex. This does not work nicely with multi-line FASTA because it will not match across the newlines. ish supports line records and also supports a generic ASCII scoring matrix. To compare accurately to agrep, we run ish in line mode with ascii scoring using a query that matches a single line of a multiline FASTA. Additionally, the scoring threshold of 0.75 for ish was translated for agrep, allowing up to 12 of the 50 characters to be errors. The same scoring of −2 for mismatch and −3 for insertions and deletions was used for both tools. The options used can be seen in Listing 5.

Ish dramatically outperforms agrep in every context. Ish is 3x faster running single threaded, scales with the number of threads, and when using both threads and GPU is 75x faster. Additionally, the ability to search FASTA records further speeds up ish, as seen in Table 2 where ish is 92x faster.

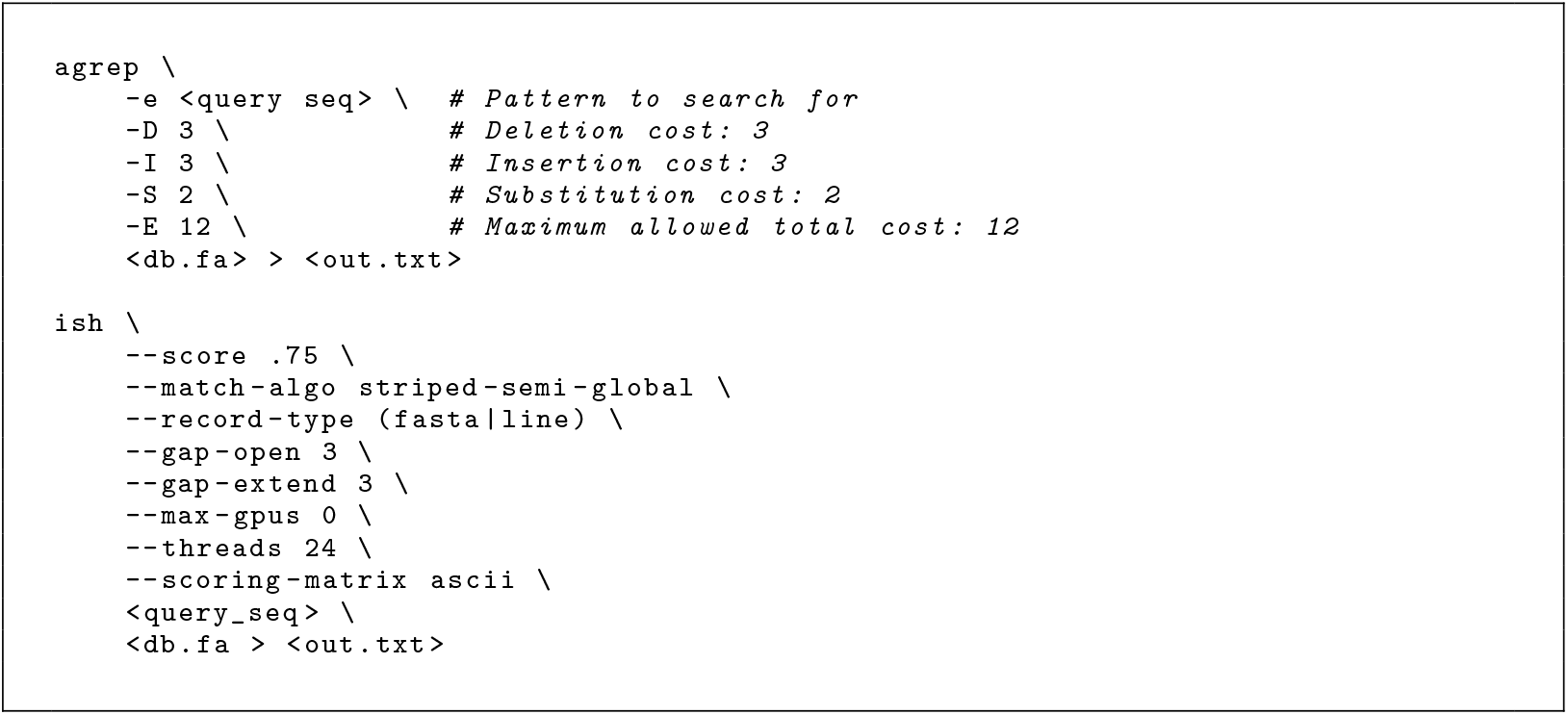
Listing 5 agrep vs ish CLI Invocations. CLI options used when running ish and agrep.

glsearch is part of the fasta36 suite of tools [26]. It does not do exactly what Ish does, but was the closest other tool found. glsearch generates alignment scores, statistics, and trace-backs for every target that is searched; it does not act as a filtering tool like ish, but more as a report generation tool like BLAST. The parameters can be seen in Listing 6 and were tuned to output something similar to ish. glsearch does glocal search, which is what ish’s semi-global is configured for by default. As seen in Table 8, ish is 1.38x faster in the best case, and even when only using the CPU dataflow, it is still 1.26x faster.

**Table 8.** Searching FASTA Records.

| Tool | Query Length | Time | Times Slower |
| --- | --- | --- | --- |
| glsearch36 | 189 | 6.756 s | 1.38 |
| ish cpu | 189 | 5.511 s | 1.12 |
| ish 1 gpu | 189 | 5.912 s | 1.21 |
| ish 2 gpu | 189 | 4.946 s | 1.01 |
| ish 3 gpu | 189 | 4.904 s | 1.00 |
| ish 4 gpu | 189 | 5.137 s | 1.05 |

Comparison of semi-global alignments between `glsearch36` and `ish`. All runs done with 24 threads. `glsearch36` parameters were tuned to make its execution as close as possible to `ish`. Times Slower is relative to the fastest time in the benchmark group.

**Table 9.** CPU Characteristics. For AWS `c7a.xlarge` and `c8g.xlarge` instances used in benchmarks.

3.3.1 Hardware and Software Versions
| Property | c7a.xlarge | c8g.xlarge |
| --- | --- | --- |
| Architecture | x64 | aarch64 |
| CPU Model | AMD EPYC 9R14 | Neoverse-V2 |
| Threads per Core | 1 | 1 |
| Base Clock (MHz) | 3699 | — |
| L1 Cache (per core) | 128 KiB (I) + 128 KiB (D) | 256 KiB (I) + 256 KiB (D) |
| L2 Cache (per core) | 1 MiB | 2 MiB |
| L3 Cache (shared) | 16 MiB | 36 MiB |
| SIMD Extensions | AVX-512 (full), AVX2, SSE4.2, FMA | SVE2 (full), NEON, i8mm, bf16 |
| Hypervisor | KVM | KVM |
| NUMA Nodes | 1 | 1 |
| Virtualization | Full | Full |
| Mitigations Applied | Spectre, SSB, IBRS | Mostly not affected |

**Table 10.** GPU Characteristics. System configuration of CPU+GPU AWS g6 series instance featuring AMD EPYC 7R13 and NVIDIA L4 used in benchmarking. CPU Cores excluded because they vary by instance type when using instances with multiple GPUs.

| Property | CPU (AMD EPYC 7R13) | GPU (NVIDIA L4) |
| --- | --- | --- |
| Architecture | x64 (Zen 3) | Ada Lovelace |
| Cores / Threads | – | 7424 CUDA cores (58 SMs × 128) |
| Base Clock (MHz) | ∼2800 | 2040 (SM), 6251 (mem) |
| L1 Cache (per core) | 32 KiB (I) + 32 KiB (D) | – |
| L2 Cache (per core) | 512 KiB | – |
| L3 Cache (shared) | 32 MiB | 48 MiB (L2 total) |
| SIMD Extensions | AVX2, FMA, SSE4.2 | Tensor Cores (TF32, BF16, INT8) |
| Memory | – | 23 GiB GDDR6 |
| Memory Bandwidth | – | ∼300 GB/s |
| Compute Capability | – | 8.9 |
| PCIe Interface | – | Gen4 ×16 (max), ×8 (active) |
| ECC Support | – | Enabled |
| CUDA / Driver Version | – | CUDA 12.4 / 550.144.03 |
| NUMA Nodes | 1 | – |
| Virtualization | KVM (full) | Pass-through |

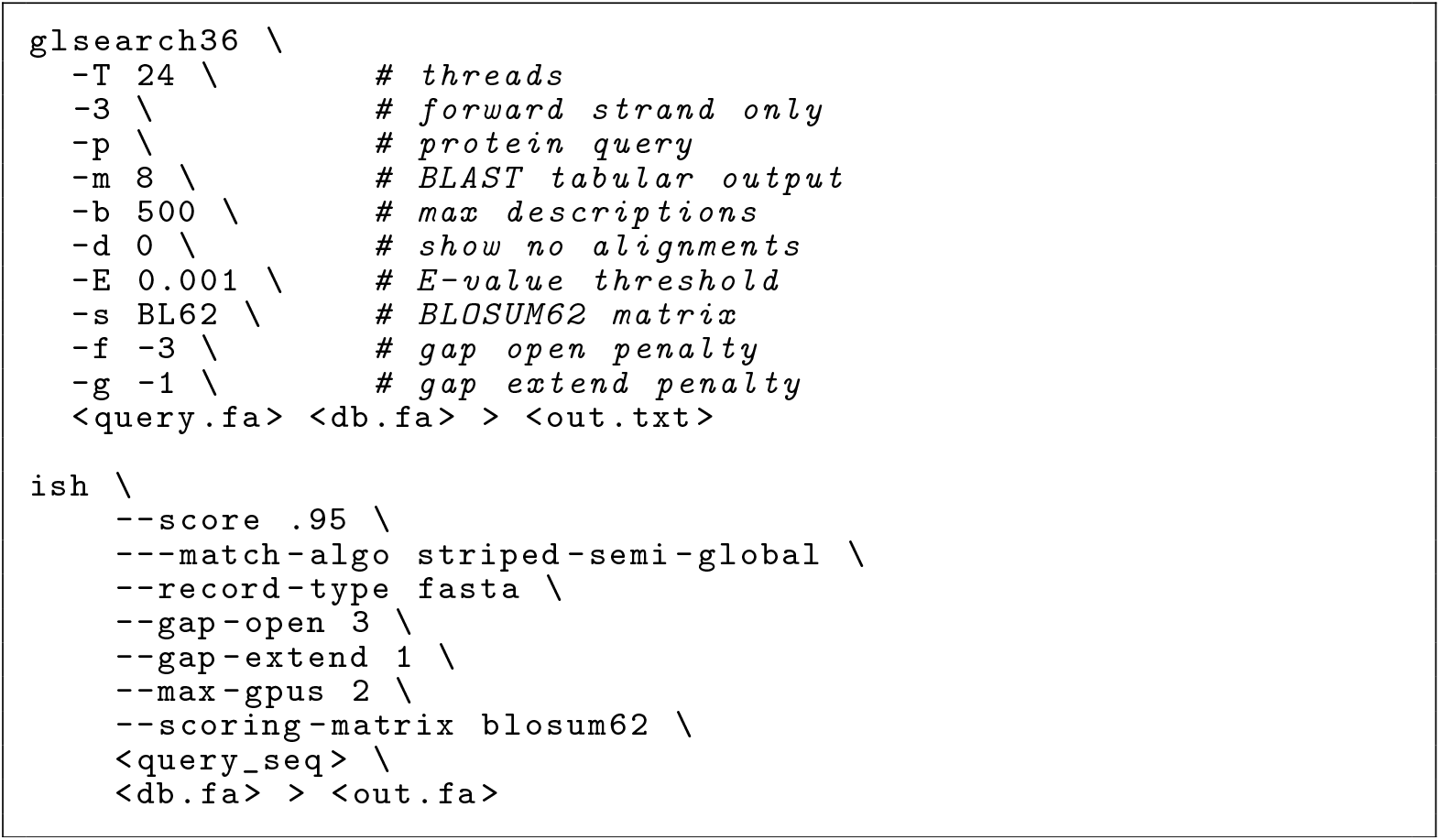
Listing 6 glsearch36 vs *ish* CLI Invocation. CLI options used when running glsearch36 and ish.

#### 3.3.1 Hardware and Software Versions

agrep version 0.9.0 was used and built out of the TRE repository [2]. glsearch version 36.3.8i as part of the fasta36 distribution [27]. Mojo v25.2.0 was used for ish [28].

## 4 Conclusions

Ish fills a niche that no other tool currently occupies; fuzzy matching on the CLI that is record type aware and uses a full alignment algorithm. This is a significant step forward from agrep [2]in terms of performance and a useful step forward from fasta36 [26] in terms of ergonomics. Ish should be useful to a wide audience for approximate matching against ASCII text. For bioinformatics, ish has a role in the development process, checking outputs for specific sequences, and possibly even as a filtering tool in pipelines to help filter input reads for specific splice sites or other unique features.

Mojo’s key strength is that it allows for creating gpu-able CLI tools without CUDA while simultaneously providing a performant SIMD version of the same algorithm. Mojo’s seamless integration with Conda makes deployment and installation incredibly easy across platforms, an issue that has plagued similar tools. Mojo’s metaprogramming capabilities set it apart in being able to leverage specifically compiled code-paths for different scenarios [29].

Using a high-level abstraction over SIMD intrinsics was effective. Performance was greater than or equal to the version that used manually written intrinsics. No regressions were observed. Additionally, the high-level abstraction allowed for novel improvements, such as the use of unsigned integers for the striped semi-global implementation. It also allows for easy use of vector strip-mining to oversubscribe SSE vectors to act more like AVX2, which would be very difficult to implement using intrinsics due to the complexity of managing cross-lane carry propagation, masking, and tail-handling logic manually across multiple SIMD registers. This could be carried forward into NEON implementations of these algorithms in C/C++/Rust for dramatic performance increases on long queries.

Ish is just scratching the surface of what is possible with GPU programming in Mojo. The algorithm is simply a low-memory scalar CPU implementation. The fact that performance that is on par with current standards can be achieved with this approach is exciting. These results demonstrate not only the immediate utility of ish, but also highlight Mojo’s potential as a platform for rapidly developing high-performance, portable sequence tools that bridge the gap between CLI usability and modern hardware acceleration.

## Declarations

### Funding

Not applicable. This work was conducted as part of my employment at Bio-Rad Laboratories.

### Conflict of interest/Competing interests

I am employed by Bio-Rad Laboratories, a for-profit company. I consulted informally with an employee at Modular Inc.—the developers of Mojo—regarding implementation details of the GPU component.

### Ethics approval and consent to participate

Not applicable.

### Consent for publication

Not applicable.

### Data availability

Not applicable.

### Materials availability

Not applicable.

### Code availability

The source code for the tool presented in this manuscript is available at: https://github.com/BioRadOpenSource/ish/. It is released under the Apache-2.0 License.

### Author contribution

I am the sole author of this work and was responsible for the conception, implementation, analysis, and writing of this manuscript.

## Notes

https://github.com/BioRadOpenSource/ish/

### Competing Interest Statement

The author is an employee of Bio-Rad Laboratories, a for-profit company. The author also had informal technical discussions with an employee of Modular Inc., the developer of the Mojo programming language used in this work. No payments or services were received from Modular Inc. or any third party in relation to this manuscript.

